# Evolutionary history of metazoan TMEM16 family

**DOI:** 10.1101/2022.04.22.489151

**Authors:** Xuye Yuan, Yu Zhu, Tatsuhiko Kadowaki

## Abstract

Transmembrane protein 16 (TMEM16) functions as either a Ca^2+^-activated Cl^-^ channel (CaCC) or phospholipid scramblase (CaPLSase) and plays diverse physiological roles. It is well conserved in eukaryotes; however, the origin and evolution of different subfamilies in Metazoa are not yet understood. To uncover the evolutionary history of the TMEM16 family, we analyzed 412 proteins from 74 invertebrate species using evolutionary genomics. We found that the TMEM16C–F and J subfamilies are vertebrate-specific, but the TMEM16A/B, G, H, and K subfamilies are ancient and present in many, but not all metazoan species. The most ancient subfamilies in Metazoa, TMEM16L and M, are only maintained in limited species. TMEM16N and O are Cnidaria- and Ecdysozoa-specific subfamilies, respectively, and Ctenophora, Xenacoelomorpha, and Rotifera contain species-specific proteins. We also identified TMEM16 genes that are closely linked together in the genome, suggesting that they have been generated via recent gene duplication. The anoctamin domain structures of invertebrate-specific TMEM16 proteins predicted by AlphaFold2 contain conserved Ca^2+^-binding motifs and permeation pathways with either narrow or wide inner gates. The inner gate distance of TMEM16 protein may have frequently switched during metazoan evolution, and thus determined the function of the protein as either CaCC or CaPLSase. These results demonstrate that TMEM16 family has evolved by gene gain and loss in metazoans, and the genes have been generally under purifying selection to maintain protein structures and physiological functions.

## Introduction

Transmembrane protein 16 (TMEM16; anoctamin, ANO) is a family of transmembrane proteins that play critical physiological functions in various cell types, such as muscle cells, neurons, and epithelial cells. TMEM16A and B are Ca^2+^-activated Cl^-^ channels (CaCCs) that are associated with epithelial fluid secretion, neuronal excitability, olfactory transduction, nociception, and photoreception (Tian, Schreiber and Kunzelmann, 2012). However, other subfamilies are not CaCCs, and most of them (TMEM16C-G, J, and K) act as Ca^2+^-activated phospholipid scramblases (CaPLSases) (Whitlock and Hartzell, 2017). CaPLSase facilitates non-specific passive phospholipid transport between the inner and outer leaflets of the membrane. For example, TMEM16F mediates the transport of phosphatidylserine from the inner to the outer leaflet of the plasma membrane, which is necessary for apoptosis and blood clotting (Suzuki *et al.*, 2010). Non-selective ion transport is also associated with phospholipid transport activity (Kostritskii and Machtens, 2021). As two fungal TMEM16 proteins, nhTMEM16 and afTMEM16, function as CaPLSases (Malvezzi *et al.*, 2013; Lee, Menon and Accardi, 2016), TMEM16A and B appear to have acquired CaCC activity during evolution.

The structures of mouse TMEM16A and F (Dang *et al.*, 2017; Paulino *et al.*, 2017; Alvadia *et al.*, 2019; Feng *et al.*, 2019) and human TMEM16K (Bushell *et al.*, 2019) together with fungal nhTMEM16 (Brunner *et al.*, 2014) and afTMEM16 (Falzone *et al.*, 2019) were determined using X-ray crystallography and/or cryogenic electron microscopy (cryo-EM). All proteins exist as homodimers with a similar overall architecture containing the specific organization of 10 transmembrane (TM) domains, suggesting that this structure is crucial for both CaCC and CaPLSase activities. Ca^2+^-binding to the major sites present in TM domains 6–8 is considered to induce the movement of TM6 at the hinge region to open the gates for TMEM16A and F proteins (Peters *et al.*, 2018; Alvadia *et al.*, 2019; Feng *et al.*, 2019). Three hydrophobic amino acids constructing the inner gate of the permeation pathway for TMEM16A and F were also identified (Lam *et al.*, 2021; Le *et al.*, 2019). According to the positions of these amino acids, the structures of mouse TMEM16A and F determined using cryo-EM appear to represent a partially opened conformation with Ca^2+^ binding. Meanwhile, the structures of TMEM16K determined using X-ray crystallography and cryo-EM are likely to represent open and closed states, respectively (Bushell *et al.*, 2019).

Due to the important physiological functions of the TMEM16 family, it is well-conserved in various eukaryotes, including plants, fungi, and animals. However, so far, research has focused on only 10 mammalian subfamilies. In fact, little is known about the functions of the invertebrate TMEM16 family. Even in fruit flies, only two proteins, Abnormal X segregation (Axs) and Subdued, have been characterized. Axs (fruit fly TMEM16K) regulates chromosome segregation and meiotic spindle assembly (Kramer and Hawley, 2003), but the CaPLSase activity has not been reported. Subdued belongs to the TMEM16A/B subfamily and is associated with host defense and thermal nociception (Wong *et al.*, 2013; Jang *et al.*, 2015). It functions as a CaPLSase (Le, Le and Yang, 2019). Furthermore, the evolution of TMEM16 genes in metazoans is not well understood. In this study, we analyzed the TMEM16 family using evolutionary genomics to provide insight into how the genes emerged and have changed in various animal species. Changes in selective pressures on TMEM16 genes during animal evolution were also studied in association with the exploitation of new ecological niches. We further compared the structures of the ANO domains of representative TMEM16 proteins predicted by AlphaFold2. Our results will serve as a foundation for the functional analyses of diverse TMEM16 proteins in future physiological and evolutionary studies.

## Results and Discussion

### Phylogeny of metazoan TMEM16 proteins

To uncover the evolutionary history of the TMEM16 family in metazoans, we collected proteins from diverse sequenced invertebrate species, together with one choanoflagellate, *Salpingoeca rosetta.* In total, 422 TMEM16 proteins, including 10 human and 12 zebrafish proteins, were phylogenetically analyzed using human transmembrane channel (TMC)-like proteins as the outgroup (Medrano-Soto *et al.*, 2018). They were classified into eight major clades (Fig. 1) with the previously identified TMEM16A/B, G, H, and K subfamilies. TMEM16L, M, N, and O are novel subfamilies specifically present in various invertebrate species.

**Figure 1.**
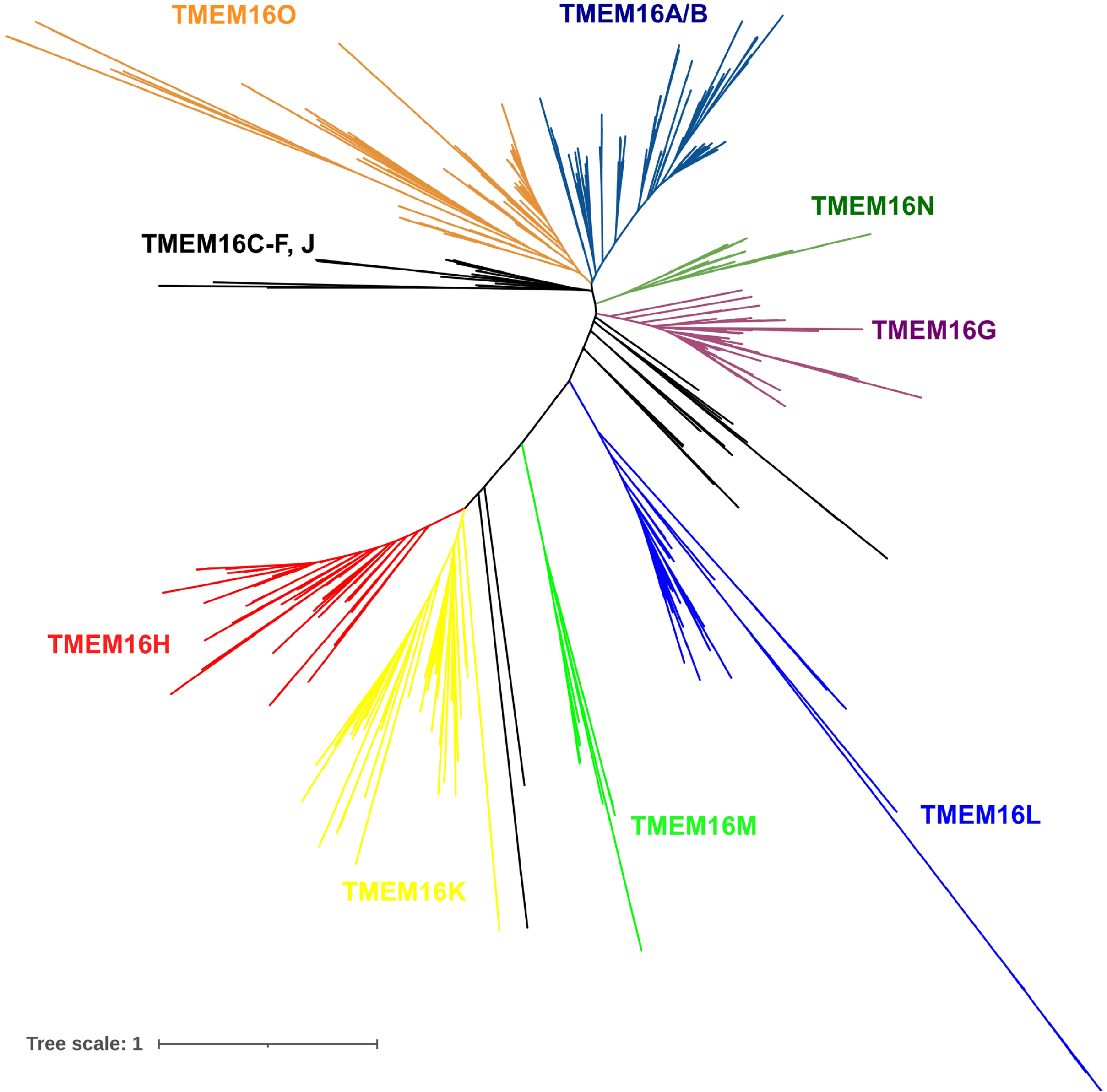
Phylogeny of the transmembrane protein 16 (TMEM16) proteins. The phylogenetic tree based on the Multiple Sequence Comparison by Log-Expectation (MUSCLE) alignment was obtained with an IQ-Tree using the LG+F+R10 best-fit model according to the Bayesian Information Criterion. The tree was unrooted, and eight human transmembrane channel-like proteins were used as outgroups. Clades corresponding to the major TMEM16 monophyletic groups are displayed in different colors. TMEM16C–F and J, as well as novel proteins that are not classified as major subfamilies, are indicated in black. Branch tags and support values are not shown to facilitate visualization, and the extended tree is presented in Supplementary Figure 1.

As previously reported (Milenkovic *et al.*, 2010), most of the invertebrates we analyzed had fewer TMEM16 proteins than vertebrates; however, Ctenophora, several Cnidaria, and *Macrostomum lignano* (Platyhelminthes) contained a larger number of proteins. Ctenophora is considered as the earliest lineage within Metazoa (Moroz *et al.*, 2014), and *Mnemiopsis leidyi* contains 14 TMEM16 proteins. There were 12 and 13 proteins in *Nematostella vectensis* and *Actinia equina* belonging to Cnidaria, respectively (Tables 1–4). These results suggest that TMEM16 genes have independently expanded in basal metazoan groups. In contrast to the 14 TMEM16 proteins in *M. lignano*, another parasitic platyhelminth, *Schistosoma mansoni,* has only four proteins, demonstrating evolutionary plasticity even in related species. Since *M. lignano* is a free-living flat worm, fewer TMEM16 proteins in *S. mansoni* could be related to parasitic life history.

**Table 1.** Composition of TMEM16 subfamilies in basal Metazoa and Deuterostomia

| Phylum | Species | A | B | A/B | C | D | E | F | G | H | J | K | L | M | N | Novel | Unassigned | Total |
| --- | --- | --- | --- | --- | --- | --- | --- | --- | --- | --- | --- | --- | --- | --- | --- | --- | --- | --- |
| Choanoflagellata | *Salpingoeca rosetta* | 0 | 0 | 0 | 0 | 0 | 0 | 0 | 0 | 0 | 0 | 0 | 0 | 0 | 0 | 4 (H/K/L/M like) | 0 | 4 |
| Ctenophora | *Mnemiopsis leidyi* | 0 | 0 | 0 | 0 | 0 | 0 | 0 | 0 | 0 | 0 | 0 | 4 | 2 | 0 | Novel: 5; K like: 1 | 2 | 14 |
| Porifera | *Amphimedon queenslandica* | 0 | 0 | 0 | 0 | 0 | 0 | 0 | 2 | 0 | 0 | 0 | 0 | 0 | 0 | 0 | 0 | 2 |
| Placozoa | *Trichoplax adhaerens* | 0 | 0 | 0 | 0 | 0 | 0 | 0 | 2 | 1 | 0 | 0 | 1 | 0 | 0 | 1 | 0 | 5 |
| Cnidaria | *Hydra vulgaris* | 0 | 0 | 0 | 0 | 0 | 0 | 0 | 3 | 0 | 0 | 1 | 1 | 1 | 4 | 0 | 0 | 10 |
|  | *Nematostella vectensis* | 0 | 0 | 1 | 0 | 0 | 0 | 0 | 2 | 1 | 0 | 1 | 1 | 1 | 5 | 0 | 0 | 12 |
|  | *Clytia hemisphaerica* | 0 | 0 | 1 | 0 | 0 | 0 | 0 | 1 | 0 | 0 | 1 | 1 | 1 | 3 | 0 | 1 | 9 |
|  | *Actinia equina* | 0 | 0 | 1 | 0 | 0 | 0 | 0 | 2 | 1 | 0 | 1 | 1 | 1 | 5 | 0 | 1 | 13 |
| Xenacoelomorpha | *Hofstenia miamia* | 0 | 0 | 0 | 0 | 0 | 0 | 0 | 3 | 0 | 0 | 0 | 0 | 1 | 0 | 4 | 1 | 9 |
| Echinodermata | *Strongylocentrotus purpuratus* | 0 | 0 | 1 | 0 | 0 | 0 | 0 | 0 | 0 | 0 | 1 | 1 | 0 | 0 | 0 | 0 | 5 |
| Hemichordata | *Saccoglossus kowalevskii* | 0 | 0 | 1 | 0 | 0 | 0 | 0 | 0 | 0 | 0 | 1 | 1 | 0 | 0 | 0 | 4 | 7 |
| Chordata | *Ciona intestinalis* | 0 | 0 | 0 | 0 | 0 | 0 | 0 | 1 | 0 | 0 | 1 | 0 | 0 | 0 | C/D/J like: 1; E/F like: 1 | 0 | 4 |
|  | *Branchiostoma lanceolatum* | 0 | 0 | 1 | 0 | 0 | 0 | 0 | 0 | 1 | 0 | 1 | 1 | 0 | 0 | 0 | 2 | 6 |
|  | *Danio rerio* | 2 | 1 | / | 1 | 0 | 2 | 1 | 1 | 1 | 1 | 2 | 0 | 0 | 0 | 0 | 0 | 12 |

“TMEM” is removed from all subfamilies for the simplicity. “Novel” represents the proteins which do not belong to the 13 major subfamilies. “Unassigned” indicates the proteins with partial ANO domains which were not phylogenetically analyzed.

**Table 2.**
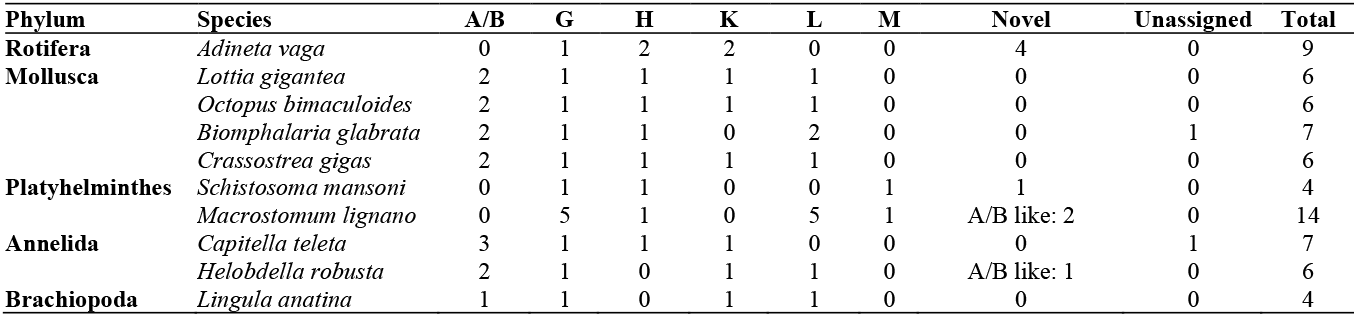
Composition of TMEM16 subfamilies in Spiralia

**Table 3.** Composition of TMEM16 subfamilies in Ecdysozoa

| Phylum/Subphylum | Species | A/B | G | H | K | L | O | Novel | Unassigned | Total |
| --- | --- | --- | --- | --- | --- | --- | --- | --- | --- | --- |
| **Hexapoda** | *Folsomia candida* | 2 | 0 | 0 | 2 | 0 | 1 |  | 0 | 5 |
| **Crustacea** | *Daphnia pulex* | 1 | 1 | 0 | 1 | 0 | 0 |  | 0 | 3 |
|  | *Penaeus vannamei* | 1 | 1 | 1 | 1 | 0 | 1 |  | 0 | 5 |
|  | *Lepeophtheirus salmonis* | 1 | 0 | 1 | 1 | 0 | 2 |  | 0 | 5 |
|  | *Tigriopus californicus* | 1 | 1 | 1 | 1 | 0 | 1 |  | 0 | 5 |
| **Myriapoda** | *Strigamia maritima* | 1 | 2 | 1 | 0 | 0 | 1 |  | 0 | 5 |
| **Chelicerata** | *Varroa destructor* | 1 | 2 | 1 | 0 | 1 | 0 |  | 0 | 5 |
|  | *Galendromus occidentalis* | 1 | 1 | 1 | 0 | 1 | 1 |  | 0 | 5 |
|  | *Dermatophagoides pteronyssinus* | 2 | 0 | 1 | 0 | 0 | 0 | H/K like: 1 | 0 | 4 |
|  | *Tetranychus urticae* | 1 | 1 | 1 | 0 | 0 | 0 |  | 0 | 3 |
|  | *Ixodes scapularis* | 1 | 1 | 1 | 0 | 1 | 1 | H/K like: 1 | 0 | 6 |
|  | *Rhipicephalus microplus* | 1 | 1 | 1 | 0 | 0 | 2 | H/K like: 1 | 1 | 7 |
|  | *Centruroides sculpturatus* | 1 | 2 | 1 | 0 | 2 | 1 |  | 1 | 8 |
|  | *Parasteatoda tepidariorum* | 1 | 1 | 1 | 1 | 2 | 1 |  | 1 | 8 |
| **Tardigrada** | *Hypsibius dujardini* | 0 | 0 | 0 | 1 | 0 | 2 |  | 1 | 4 |
|  | *Ramazzottius varieornatus* | 0 | 0 | 0 | 2 | 0 | 3 |  | 0 | 5 |
| **Nematoda** | *Caenorhabditis elegans* | 0 | 0 | 1 | 0 | 1 | 0 |  | 0 | 2 |

Tardigrada and Nematoda represent phylum and others are subphylum.

**Table 4.**
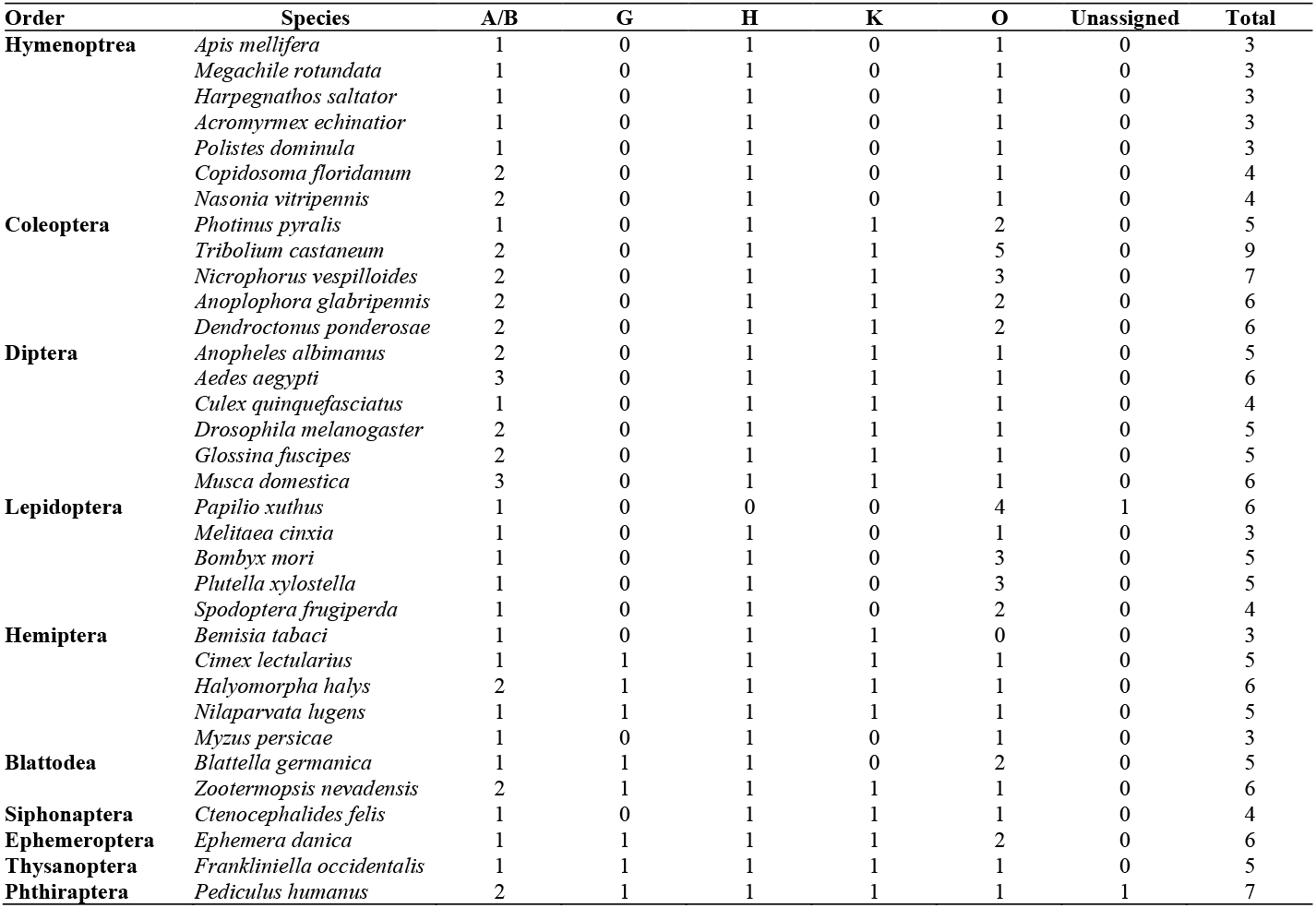
Composition of TMEM16 subfamilies in insects

### TMEM16A/B (ANO1/2) subfamily

TMEM16A and B are vertebrate-specific; however, many animal species have TMEM16A/B, which is derived from the same ancestral gene (Supplementary Fig. 1). Because this subfamily appears to be absent in Choanoflagellates, Ctenophora, Porifera, and Placozoa, the gene family was likely to emerge in the common ancestor of Eumetazoa (Cnidaria and Bilateria). It is present in most animal species, but has been lost in Tardigrada, Nematoda, several species of Spiralia, *Hydra vulgaris, Hofstenia miamia,* and *Ciona intestinalis* (Tables 1–4).

In *Capitella teleta,* two closely related TMEM16A/B genes, CapteP153854 (CAPTEscaffold_249: 186,019-195,820) and CapteP153860 (CAPTEscaffold_249: 231,455-238,517), exist in the same genomic contig approximately 35.6 kb away from each other. LOC101898447 (NW_004766068.1: 5,507-20,539) and LOC101898846 (NW_004766068.1: 32,428-41,620) are present continuously in the same contig in *Musca domestica*. The same is true for LOC5576066 (Chr1: 59,405,066-59,531,372) and LOC5576072 (Chr1: 59,556,793-59,593,868) in *Aedes aegypti*. These results suggest that TMEM16A/B gene has recently been duplicated in these three species.

### TMEM16C–F and J (ANO3–6 and 9) subfamilies

The TMEM16C-F and J subfamilies appear to be vertebrate-specific and absent in the other species (Tables 1–4). TMEM16C, D, J, and *C. intestinalis* XP 026692266.1 share the common ancestral gene. The same applies to TMEM16E, F, and *C. intestinalis* XP 026691398.1 genes (Supplementary Fig. 1). These results suggest that two ancestral genes of the TMEM16C–F and J subfamilies specifically arose in the Chordata lineage.

### TMEM16G, H, and K (ANO7, 8, and 10) subfamilies

TMEM16G and H subfamilies are present in Porifera and Placozoa, respectively. Thus, TMEM16G arose in the common ancestor of Porifera and the rest of the animal species. TMEM16H emerged in the common ancestor of Placozoa and Eumetazoa. The TMEM16K subfamily appeared to have emerged in the common ancestor of Eumetazoa. However, they were lost in many other animal species (Tables 1–4). Interestingly, TMEM16G was lost in holometabolous insects, while TMEM16H was lost in Hymenoptera and Lepidoptera (Table 4).

Recently duplicated TMEM16G genes were identified in some species. *Amphimedon queenslandica* contains LOC100640826 (NW_003546253.1: 413,296-419,184) and LOC100640950 (NW_003546253.1: 419,407-428,220) as the duplicated genes. *Trichoplax adhaerens* has duplicated genes, TRIADDRAFT_10718 (NW_002060948.1: 5,135,884-5,143,578) and TRIADDRAFT_26725 (NW_002060948.1: 5,154,343-5,159,159). In *H. vulgaris,* three genes, LOC100204318 (NW_004166878.1: 118,761-163,326), LOC100199510 (NW_004166878.1: 186,536-240,936), and LOC100203187 (NW_004166878.1: 254,783-312,418), are tandemly present in the genome. LOC5522289 (NW_001834413.1: 726,041-747,337) and LOC5522290 (NW_001834413.1: 748,312-772,062) genes locate next to each other in *N. vectensis* genome. *A. equina* contains EGACTEQ4350024001 (WHPX01000439.1: 17,082-32,783) and EGACTEQ4350037435 (WHPX01000439.1: 33,358-46,883) genes in tandem. TMEM16G was likely to duplicate in the common ancestor of Cnidaria. Among the five TMEM16G genes in *M. lignano*, PAA80633.1, PAA80632.1, and PAA80631.1, exist in the same contig, LFJF01009900.1.

### TMEM16L (ANO11) subfamily

The presence of the TMEM16L subfamily in Ctenophora suggests that it arose from the common ancestor of all animal species. However, it has been lost in many animal species, such as vertebrates and insects. Among Ecdysozoa, several species in Chelicerata and Nematoda have maintained this subfamily (Tables 1–4).

Among the four TMEM16L genes in *M. leidyi*, ML032912a (ML0329: 65,981-68,991) and ML032913a (ML0329: 69,185-72,941) were found to be recently duplicated genes. Two recently duplicated genes, LOC106075397 (NW_013193578.1: 84,919-120,292) and LOC106075340 (NW_013193578.1: 67,771-145,736), are present in the *Biomphalaria glabrata* genome. Among the five TMEM16L genes in *M. lignano*, PAA61603.1, PAA66696.1, and PAA54271.1 are present in the same contig, scaf1542, suggesting a recent gene duplication event. There are also two recently duplicated genes, LOC111623019 (NW_019384530.1: 324,628-339,902) and LOC111622994 (NW_019384530.1: 343,753-350,203) in *Centruroides sculpturatus* genome. These results suggest that duplication of TMEM16L gene occurred sporadically in a few species.

### TMEM16M (ANO12) subfamily

The TMEM16M subfamily is present in Ctenophora, *M. leidyi*, suggesting that it emerged in the common ancestor of all animal species, together with TMEM16L. However, it has been lost in many animal species and currently exists only in Ctenophora, Cnidaria, Xenacoelomorpha, and Platyhelminthes (Tables 1–4).

### TMEM16N (ANO13) subfamily

TMEM16N is a Cnidaria-specific subfamily with multiple genes (3-5) in each species. In fact, *N. vectensis* has the five genes and LOC5520273 (NW_001834404.1: 1,926,626-1,945,283), LOC5520274 (NW_001834404.1: 1,951,305-1,960,155), and LOC5520189 (NW_001834404.1: 1,961,734-1,969,704) genes are linked in the genome. In *H. vulgaris,* LOC100205448 (NW_004167726.1: 926-38,967) and LOC100210722 (NW_004167726.1: 92,469-130,728) genes are linked in the genome among the four genes. *A. equina* contains EGACTEQ4350028189-PA (WHPX01000871.1: 667,440-671,454) and EGACTEQ4350040606-PA (WHPX01000871.1: 673,569-685,782) genes by tandem in the genome among the five genes. However, no linkage was found between three TMEM16N genes in *Clytia hemisphaerica*. These results suggest that TMEM16N has undergone duplication in the common ancestor of Cnidaria, as well as in each species.

### TMEM16O (ANO14) subfamily

TMEM16O is an Ecdysozoa-specific subfamily that is absent in only a few species, such as the Nematoda. There are four clades within the subfamily, and the second largest includes multiple genes in Coleoptera and Lepidoptera. Insects of these two orders also have additional genes in the largest clade (Table 4 and Supplementary Fig. 1). We first characterized the genomic localization of genes in the second-largest clade. Two of the four *Tribolium castaneum* genes, LOC100141597 (ChrLG6: 5,266,955-5,278,052) and LOC658673 (ChrLG6: 5,284,248-5,295,823), are present in tandem in the genome. The LOC108562319 (NW_017098395.1: 2,623-5,903) and LOC108562321 (NW_017098395.1: 6,323-9,275) genes are linked together in the genome of *Nicrophorus vespilloides. Bombyx mori* LOC101739925 (Chr15: 7,612,667-7,635,5749) and LOC101739691 (Chr15: 7,636,133-7,653,974) are present next to each other in the genome. The same is true for LOC105380640 (NW_024010847.1: 11,184,657-11,276,385) and LOC105380711 (NW_024010847.1: 11,605,103-11,638,232) genes in *Plutella xylostella.* Three genes in *Papilio xuthus,* LOC106117307 (NW_013530969.1: 3,441,514-3,452,888), LOC106117111 (NW_013530969.1: 3,471,245-3,484,096), and LOC106117110 (NW_013530969.1: 3,484,310-3,505,524) are continuously located in the same genomic contig. However, three Coleopteran insects, *Photinus pyralis, Anoplophora glabripennis, Dendroctonus ponderosae* have single genes and two Lepidopteran insects, *Melitaea cinxia* and *Spodoptera frugiperda* do not contain the gene in this clade. Furthermore, two hemimetabolous insects, *Blattella germanica* and *Ephemera danica* also have a single gene. These results suggest that the ancestral gene of TMEM16O duplicated once and that one copy has been lost in most insects, but has undergone recent duplication in the above species of Coleoptera and Lepidoptera. However, it is also possible that gene duplication occurred in the common ancestor of holometabolous insects, followed by the loss of one copy in Hymenoptera, Diptera, and some Coleoptera and Lepidoptera species.

Both *Ramazzottius varieornatus* and *Hypsibius dujardini* have multiple genes in the Tardigrada-specific clade (Supplementary Fig. 1). Among the three genes in *R. varieornatus,* RVY_01825 (BDGG01000001.1: 5,236,110-5,241,178) and RvY_01836 (BDGG01000001.1: 5,266,086-5,269,655) are tandemly present in the genome. Two *H. dujardini* genes, OQV17298.1 (MTYJ01000062: 42736-35420) and OQV17300.1 (MTYJ01000062: 54029-44731), are located in the same genomic contig. Thus, gene duplications have also recently occurred in the Tardigrada lineage. Finally, two TMEM16O genes in *Spodoptera frugiperda* appear to have been generated by a recent duplication, as LOC118280504 (Chr29: 9,494,944-9,524,103) and LOC118280221 (Chr29: 9,529,882-9,586,765) are present in tandem in the genome. The estimated time for the birth of each TMEM16A/B–O subfamily member during metazoan evolution is summarized in Figure 2. They have not been stable and have undergone complex patterns of gene loss and duplication, as described above (Tables 1–4).

**Figure 2.**
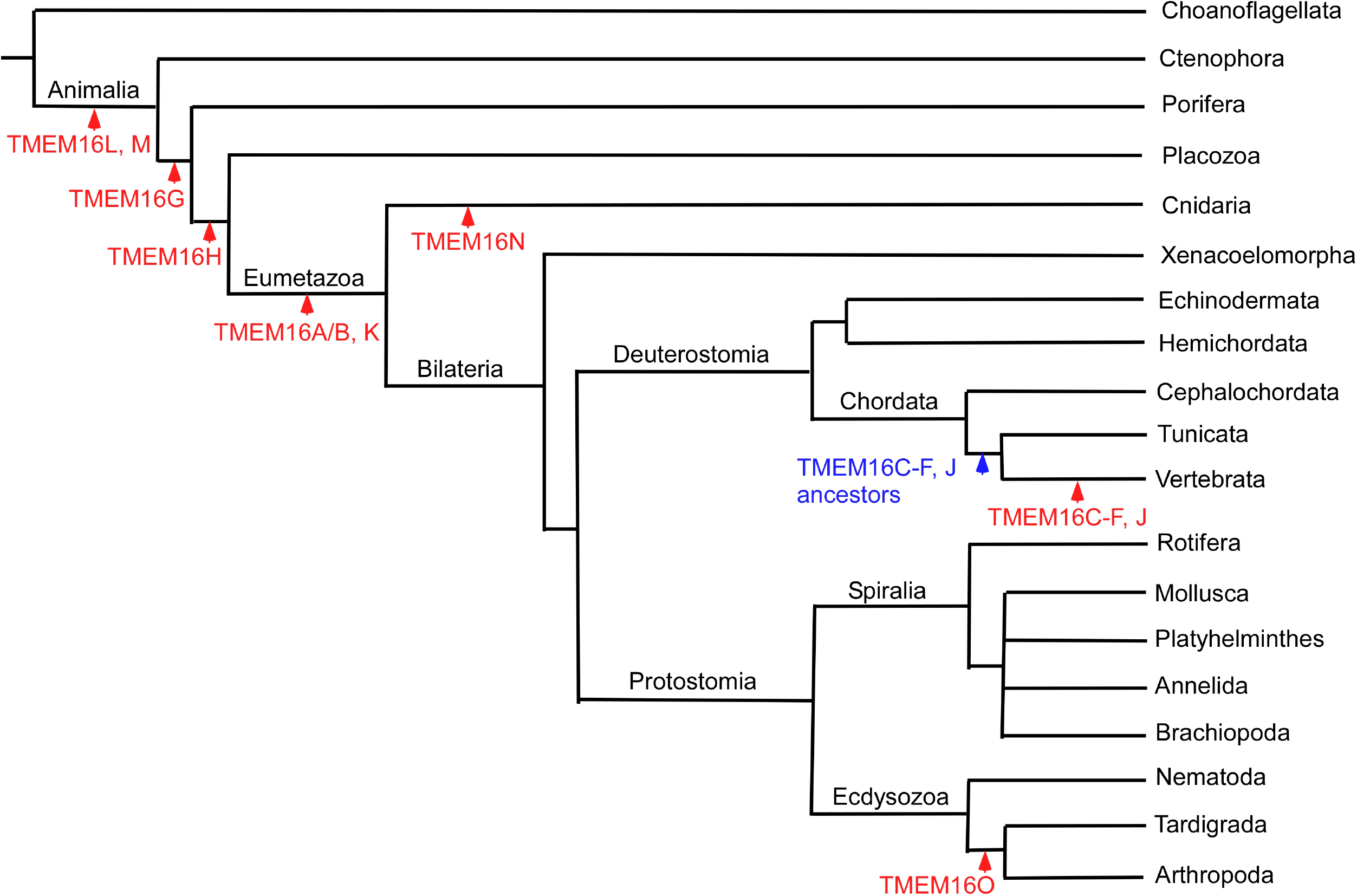
A model for the emergence of TMEM16 genes. Schematic phylogenetic tree highlighting the branches along which specific TMEM16 subfamily genes appeared (red). They were inferred from their presence or absence in the tested genomes of the extant species. The branch lengths contain no temporal information. It is possible that the TMEM16O subfamily appeared in the ancestor of Ecdysozoa followed by the loss in Nematoda. Two ancestral genes for the TMEM16C–F and J subfamilies (blue) might exist in the ancestor of Chordata as there are two unassigned TMEM16 proteins in *Branchiostoma lanceolatum* (Table 1). After emergence, each TMEM16 gene underwent complex patterns of duplication and/or loss (Tables 1–4).

### Novel TMEM16 proteins

Our analysis revealed TMEM16 proteins that cannot be classified into the above eight major subfamilies. *S. rosetta* has four TMEM16 genes that share a common ancestor with TMEM16H, K, L, and M. *Adineta vaga* and *M. leidyi* also contains four and five genes which would be related to *S. rosetta*’s genes, respectively (Supplementary Fig. 1). There are four novel TMEM16 proteins that are distinct from all other proteins in *H. miamia* (Supplementary Fig. 1). These results suggest that TMEM16 can expand independently in individual species.

### ANO domain structures of TMEM16A/B and L–O proteins

To provide insights into the structure-function relationships of invertebrate-specific TMEM16A/B and L-O proteins, we first examined the protein domains. All subfamilies, except TMEM16M, have both an ANO domain (pfam04547) and an ANO dimerization domain (pfam16178). TMEM16M proteins are relatively small compared to other subfamily proteins, and only *N. vectensis* and *A. equina* proteins have a partial ANO dimerization domain. Since human TMEM16K without this domain was shown to form a homodimer (Bushell *et al.*, 2019), it is not necessary for TMEM16 dimerization. For further analyses, we selected a single *A. equina* protein for each TMEM16A/B and L–N subfamily, and *Apis mellifera* TMEM16O as representatives. We aligned the amino acid sequences and particularly focused on TM3-8 domain sequences, which constitute the permeation pathway, as well as the Ca^2+^ binding site in TMEM16A, F, and K proteins (Fig. 3). Five Glu/Asp residues responsible for Ca^2+^ binding in the TM6–8 domains are well-conserved, suggesting that all proteins are activated by intracellular Ca^2+^. Among the five hydrophobic amino acids that function as the inner gates (at the neck regions) for the pores of TMEM16A and F proteins, two Ile residues in the TM4 and 6 domains (corresponding to I550 and I641 of TMEM16A) are conserved and others are partially conserved. Although AeTMEM16L and AmTMEM16O have Met residues at a site equivalent to I551 of TMEM16A, 13 TMEM16L and 26 TMEM16O proteins have hydrophobic amino acids. Thus, most TMEM16A/B and L–O proteins are likely to have similar inner gate structures. The Gly residue at the hinge region of the TM6 domain is also conserved, except for the TMEM16L subfamily, in which only two *M. lignano* proteins (PAA61603.1 and PAA66696.1) contain Gly residues.

**Figure 3.**
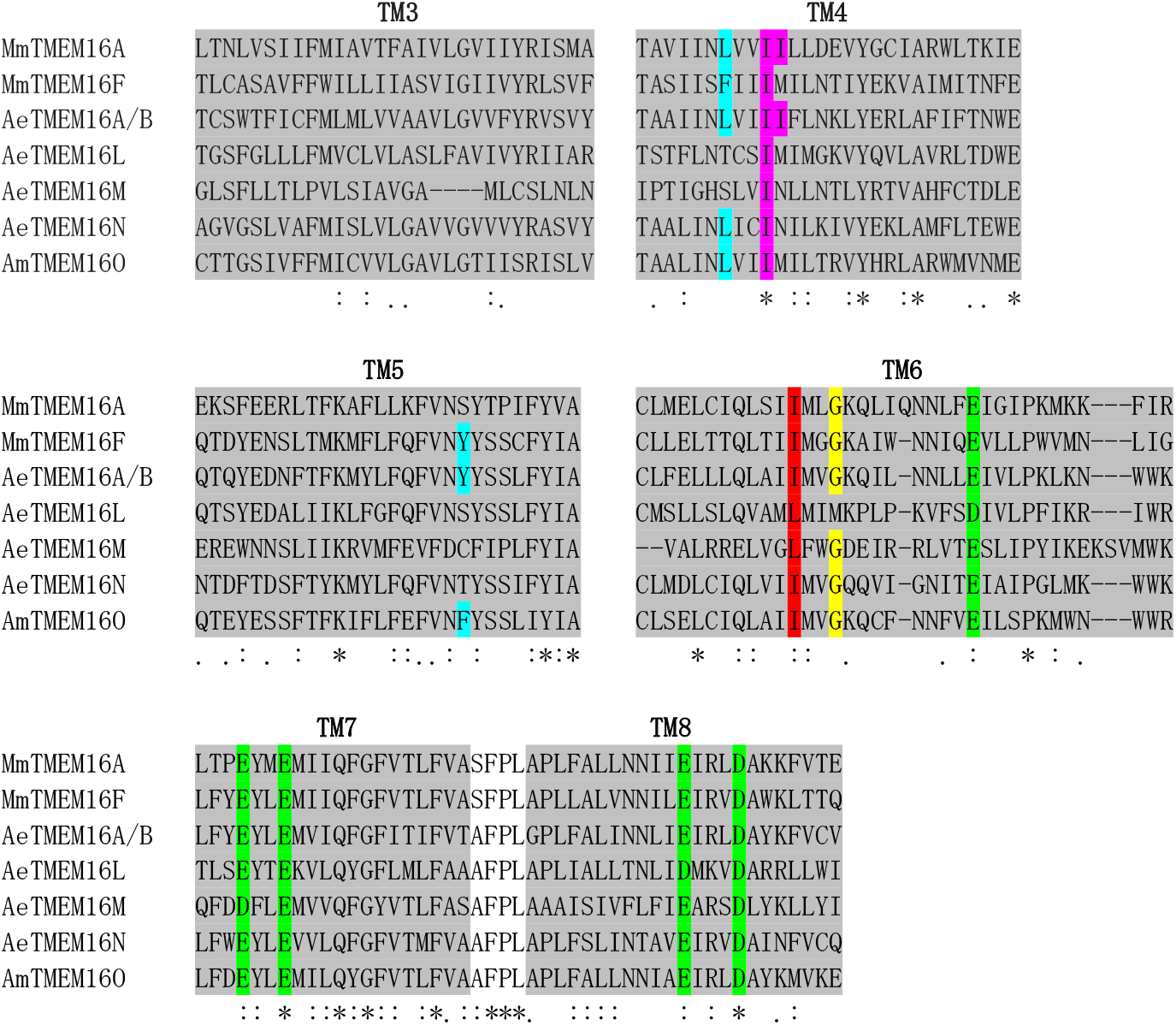
Sequence analysis of the transmembrane (TM)3-8 domains in TMEM16A, F, A/B, L-N, and O. Amino acid sequences of murine (Mm) TMEM16A and F, beadlet anemone (Ae) TMEM16A/B, L-N, and honey bee (Am) TMEM16O were aligned using Multiple Alignment using Fast Fourier Transform (MAFFT), and the TM3–8 domain sequences are highlighted in light gray. The hydrophobic amino acid residues corresponding to the inner gate of TMEM16A (I550 and I551) are highlighted in purple. The hydrophobic amino acid residues corresponding to the inner gate of TMEM16F (F518 and Y563) are highlighted in light blue. Hydrophobic amino acid residues corresponding to the third residue of the inner gates of TMEM16A and F (TMEM16A: I641; TMEM16F: I612) are shown in red. Gly residues at the hinge regions of TM6 domains are highlighted in yellow. All proteins contain highly conserved Ca^2+^-binding residues in the TM6, 7, and 8 domains (highlighted in green).

We also predicted the ANO domain structures of the TMEM16A/B and L–O proteins using AlphaFold 2 (Jumper *et al.*, 2021). As shown in Fig. 4, the overall architectures of the proteins were similar, with the same arrangement of 10 TM domains. Five putative Glu/Asp residues necessary for Ca^2+^ binding are also clustered below the hinge region of the TM6 domain in all the proteins (Fig. 4A, C, E, G, and I). We measured the distance between two well-conserved hydrophobic amino acids that form the inner gate in the TM4 and 6 domains. As a reference, the ANO domain structures of mouse TMEM16A and F were determined using AlphaFold2 (Supplementary Fig. 2A and B). TMEM16A and AeTMEM16A/B have narrow inner gates with 4.2 and 4.7 Å, respectively. Meanwhile, TMEM16F, AeTMEM16L, -M, -N, and AmTMEM16O have wide inner gates with 9.9, 8.6, 8.8, 8.5, and 9.1 Å, respectively (Fig. 4B, D, F, H, and J). The distances of the narrow and wide inner gates correspond well to those of TMEM16K in the closed and open states, respectively (Bushell *et al.*, 2019). The human TMEM16K structure predicted by AlphaFold2 contains an inner gate of 9.3 Å (Supplementary Fig. 2C), which is equivalent to that with the opened conformation determined by X-ray crystallography (PDB ID: 6R65 and 5OC9). These results suggest that the function of TMEM16 protein as either CaCC or CaPLSase can be inferred by the distance of the inner gate made by TM4 and TM6 domains using AlphaFold2. The extracellular portion of the TM4 domain localizes in front of the TM5 domain to close the groove and construct the pore necessary for ion permeation in TMEM16A (Falzone *et al.*, 2018). Therefore, we determined the ANO domain structures of other TMEM16A/B proteins. In contrast to AeTMEM16A/B, the distances of inner gates in two Cnidarian orthologs, NvTMEM16A/B and ChTMEM16A/B, are 9.2 and 9.7 Å, respectively (Supplementary Fig. 3A–C). These results indicate that the inner gate width can vary between orthologs in the same phylogenetic clade. Each species in Mollusca and Annelida contains 2-3 TMEM16A/B proteins which are clustered to the different clades except *Helobdella robusta* having two proteins in a single clade (Supplementary Fig. 1 and Table 2). *Lottia gigantea* protein, XP.009060312.1, and two *C. teleta* proteins, CapteP153854 and CapteP153860, are grouped in the same clade (Supplementary Fig. 1) but had inner gates of 9.6, 5.0, and 4.5 Å, respectively. Meanwhile, the inner gate distances in *L. gigantea* protein, XP.009065298.1 and *C. teleta* protein, CapteP222303 belonging to the other clade are 4.6 and 8.6 Å, respectively (Supplementary Fig. 3D–H). Similarly, most Coleoptera and Diptera also contain 2–3 TMEM16A/B proteins classified into different clades (Supplementary Fig. 1 and Table 4). The distances of inner gates in *T. castaneum* protein (XP.008192276.1) and *Drosophila melanogaster* NP.001245633.1 (CG10353) grouped in one clade are 4.7 and 4.6 Å, respectively. *T. castaneum* protein (XP.008193545.1) and *D. melanogaster* NP.001189248.1 (Subdued) in the other clade have the inner gates with 9.3 and 8.9 Å, respectively (Supplementary Fig. 3I–L). Subdued is a CaPLSase (Le, Le and Yang, 2019) but the function of CG10353 remains to be determined. The above results suggest that the inner gate distance varies between orthologous TMEM16A/B proteins in the same phylogenetic clade; however, the proteins classified into different clades in a single species have either narrow or wide gates. TMEM16 protein may have frequently switched the function as either CaCC or CaPLSase during metazoan evolution. These results are also consistent with the fact that TMEM16A can be converted to CaPLSase by a single amino acid substitution (L547K, K588N, and V543S) (Jiang *et al.*, 2017; Le *et al.*, 2019), but the inner gate widths of these mutant proteins remain to be determined.

**Figure 4.**
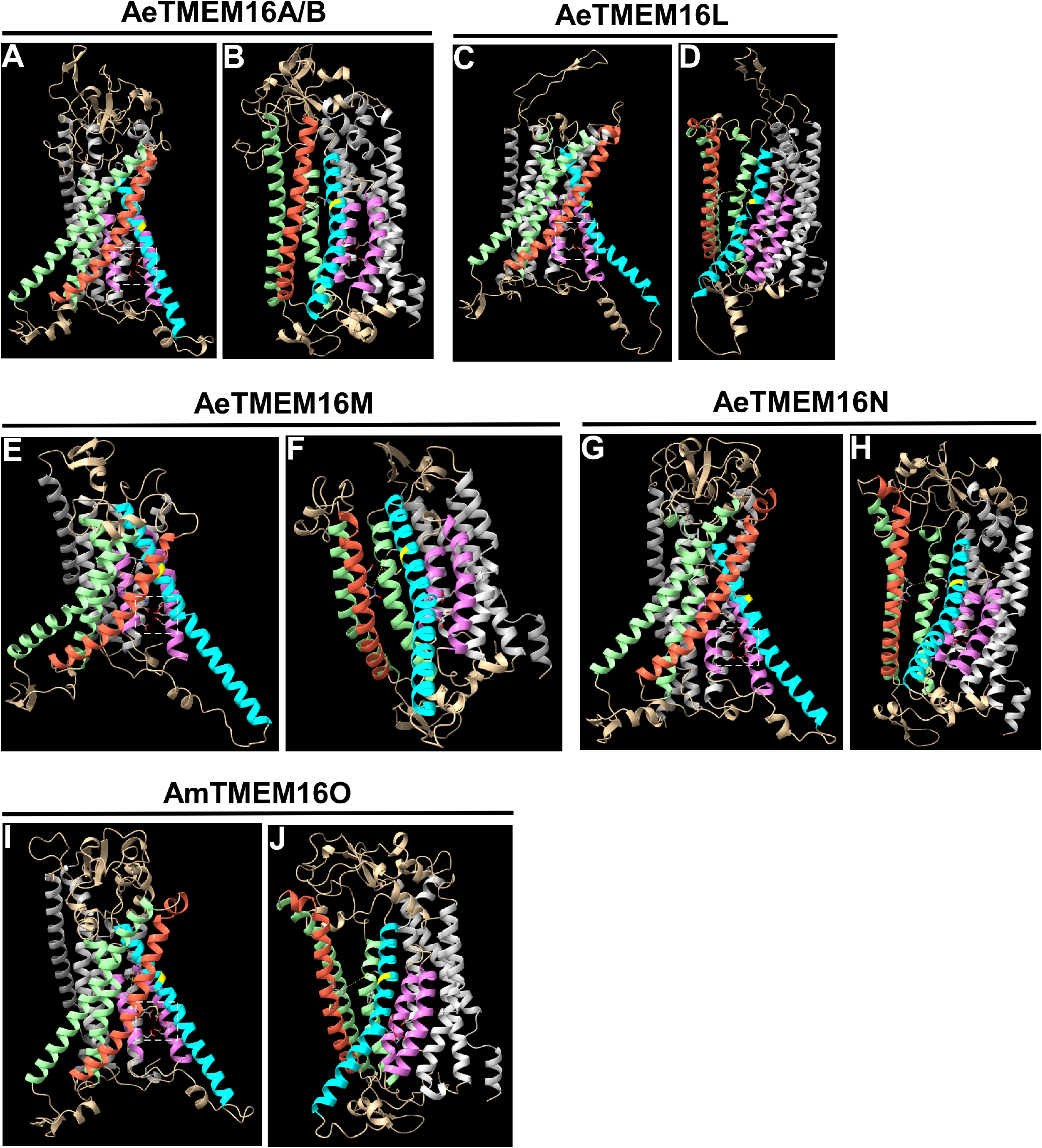
Anoctamin (ANO) domain structures of TMEM16A/B, L-N, and O. Overall structures of ANO domains in AeTMEM16A/B, L-N, and AmTMEM16O predicted by AlphaFold2 are shown. The TM1, 2, 9, and 10 domains are shown in silver, while the TM3 and 5 domains are highlighted in pale green. TM7 and 8 domains are highlighted with orchid, while the TM4 and 6 domains are shown in tomato and cyan, respectively. Five putative Ca^2+^-binding Glu/Asp residues in the TM6, 7, and 8 domains are indicated by squares with white dotted lines (A, C, E, G, and I). The side chains of the five potential amino acids constructing the inner gate in the TM4, 5, and 6 domains are also shown. The Gly residue (or Met for AeTMEM16L) in the main chain is highlighted in yellow at the potential hinge region of the TM6 domain. The distance between two hydrophobic amino acid residues in the TM4 and 6 domains to form the inner gate (corresponding to I550 and I641 of TMEM16A) is also indicated (yellow dotted line) (B, D, F, H, and J).

### Selective pressure on TMEM16 genes

We analyzed the selective forces acting on TMEM16A/B, G, H, and K–O subfamilies by calculating the ratio of non-synonymous to synonymous nucleotide substitution rates (ω, *d*_N_/*d*_S_) for all tested species. Most amino acid residues of TMEM16 proteins are expected to have evolved under purifying selection to maintain conserved structural and functional properties. However, there may be a small number of positively-selected amino acids that contribute to their functional diversity. All the tested genes have evolved under purifying selection (ω « 1), revealing that all of them encode functional proteins. TMEM16H and M subfamilies show the lowest *d*_N_/*d*_S_ ratio, suggesting that they may have very conserved roles in various species. The TMEM16N subfamily had the highest *d*_N_/*d*_S_ ratio (0.07), suggesting that it has evolved under weaker purifying selection and/or contains more sites that have been shaped by positive selection (Table 5). This is consistent with the fact that TMEM16N is a Cnidaria-specific subfamily that contains recently duplicated genes. Site-specific positive selection may be more easily detectable in recently duplicated TMEM16 genes that potentially undergo functional divergence. Thus, we calculated the *d*_N_/*d*_S_ ratios for these TMEM16 genes. As shown in Table 5, some of these gene sets had higher *d*_N_/*d*_S_ ratios than those calculated using all the genes in each subfamily. The Lepidoptera TMEM16O genes in the second-largest clade have the highest *d*_N_/*d*_S_ ratio (0.18). We applied the site class models, M7 and M8, in phylogenetic analysis by maximum likelihood method to obtain evidence for site-specific selection. However, positively-selected sites with a posterior probability of > 95% were not identified. Thus, it is likely that the evolution of the TMEM16 family has been driven by gene gain and loss rather than the positive selection of specific amino acids during metazoan diversification.

**Table 5.**
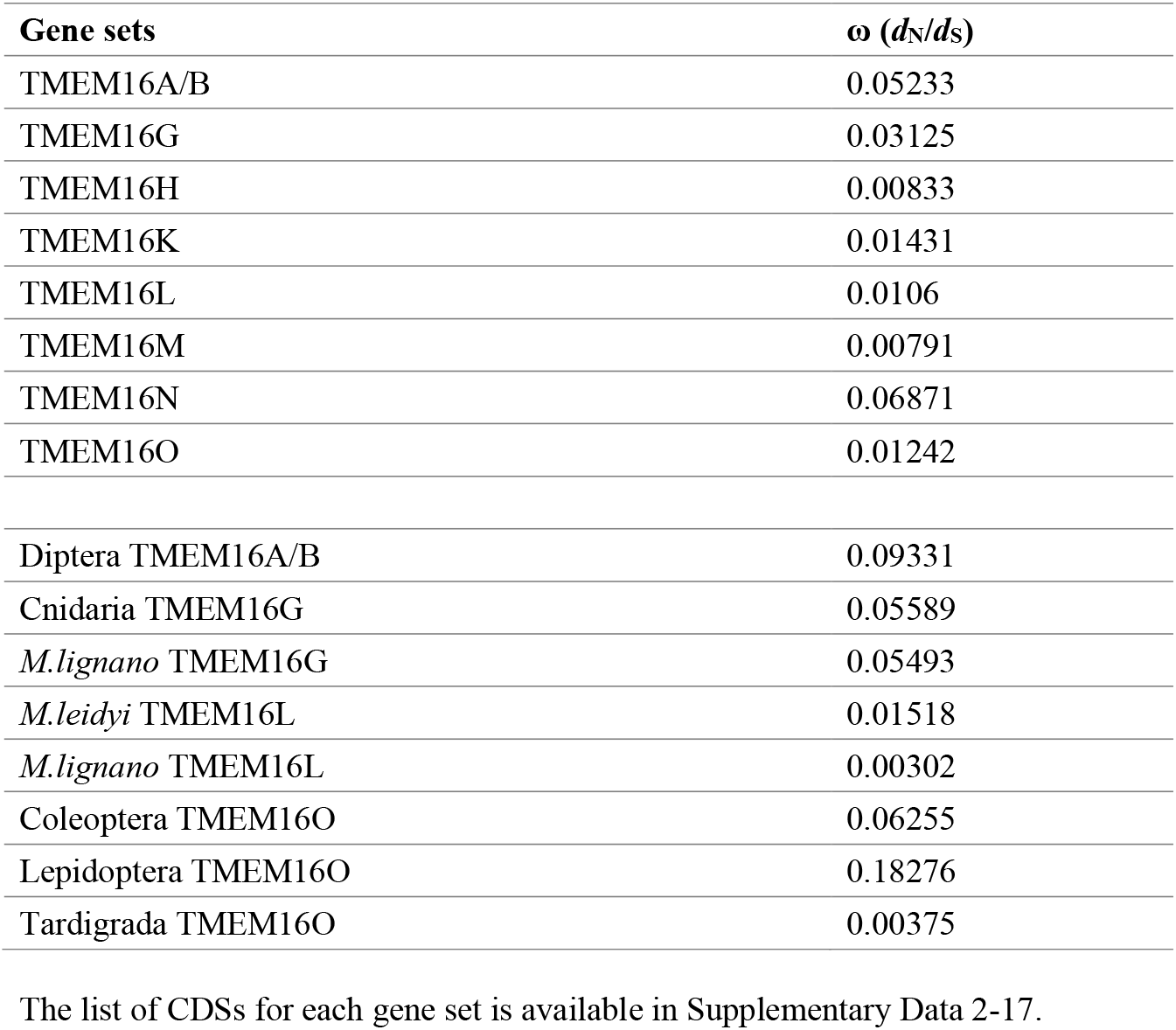
Selective pressures on TMEM16 genes

## Materials and Methods

### Collection of TMEM16 proteins

We first identified eight *T. castaneum* TMEM16 proteins from the annotated protein sets. These red flour beetle and 10 human TMEM16 protein sequences were used as the queries to search for TMEM16 proteins in the organisms listed in Tables 1–4 by BLASTP search with the cut-off E-value of 1E-10 against annotated protein databases. We then analyzed all of the collected proteins by Conserved Domains search (https://www.ncbi.nlm.nih.gov/Structure/bwrpsb/bwrpsb.cgi) to confirm they contain the full length ANO domain (pfam04547). The proteins with the partial ANO domains were not subjected to the phylogenetic analysis and classified as “unassigned” in Tables 1–4. 400 TMEM16 proteins were selected and they were analyzed for the phylogenetic tree together with 10 human and 12 zebrafish TMEM16 proteins. The analyzed sequences are presented in Supplementary Data 1. The protein databases used were NCBI (http://www.ncbi.nlm.nih.gov), EnsemblMetazoa (http://metazoa.ensembl.org/index.html), EnsemblProtists (http://protists.ensembl.org/index.html), and InsectBase (http://v2.insect-genome.com/).

### Phylogenetic analysis of TMEM16 proteins

We used MUSCLE v5 program (Edgar, 2021) to align the amino acid sequences of above 422 TMEM16 proteins and eight human TMC-like proteins as the outgroup. The aligned sequence was then trimmed using trimA1 tool (Capella-Gutiérrez, Silla-Martínez and Gabaldón, 2009). Following the trimming, we built a phylogenetic tree via maximum likelihood method using IQ-TREE 2 (Minh *et al.*, 2020). The best-fit amino acid substitution models were selected by applying the ModelFinder program (Kalyaanamoorthy *et al.*, 2017), and LG+F+R10 was chosen according to Bayesian information criterion. The bootstrap value was estimated using ultrafast bootstrap approximation (Minh, Nguyen and von Haeseler, 2013; Hoang *et al.*, 2018) with 10,000 replications in IQ-TREE 2. We viewed and graphically edited the tree with iTOL (Letunic and Bork, 2021).

### Locations of phylogenetically related TMEM16 genes on the genomes

Based on the phylogenetic tree of 422 TMEM16 proteins, we identified many sets of the closely related proteins in single species. The genes encoding these proteins were analyzed by Genome Data Viewer in NCBI (https://www.ncbi.nlm.nih.gov/genome/gdv/?org) and EnsemblMetazoa to determine their locations in the genome. In case they are closely linked, the results are indicated by the positions in the specific chromosome or contig. Because a genome browser is not available for *M. lignano*, we analyzed the genomic contig sequences by TBLASTN using the candidate protein sequences as the queries.

### Amino acid sequence alignment

We aligned the amino acid sequences of murine TMEM16A and F, beadlet anemone TMEM16A/B, L-N, as well as honey bee TMEM16O using MAFFT (https://www.ebi.ac.uk/Tools/msa/mafft/). *A. equina* contains five TMEM16N proteins (Table 1) and utg935.6 peptide was used for the alignment as the representative. We identified and analyzed TM3-8 domains to compare the amino acid residues for potential inner gate, hinge region of TM6 domain, and Ca^2+^ binding.

### Structural determination using AlphaFold2

We first identified the amino acid sequence representing ANO domain in each target protein using Conserved Domains search. The structures of these ANO domains were then determined using ColabFold (Mirdita *et al.*, 2021) based on AlphaFold v2.1.0 (Jumper *et al.*, 2021). For human TMEM16K, the full length sequence was used. The generated structures were analyzed and displayed using UCSF ChimeraX (Pettersen *et al.*, 2021; Goddard *et al.,* 2018).

### Analysis of the selection pressures on TMEM16 genes

We first obtained the multiple alignment of amino acid sequences (MSA) with MUSCLE and used it to guide the corresponding CDS sequence alignments with PAL2NAL (Suyama, Torrents and Bork, 2006). We then built the phylogenetic trees from the MSA after trimming with trimA1 tool. CodeML program in the PAML v4.9 package (Yang, 2007) was used to estimate synonymous substitution rate ω (ω = *d*_N_/*d*_S_ = nonsynonymous/synonymous substitutions) with the CDS sequence alignments and phylogenetic trees as inputs for positive selection. The neutral model M0 was used to obtain ω, while ω > 1 indicated positive selection. We compared null model M7 and positive selection model M8 using Likelihood Ratio Test with two degrees of freedom (Yang *et al.*, 2000) and allowed the rejection of the null model (P < 0.05). The Bayes Empirical Bayes approach (Yang, Wong and Nielsen, 2005) with posterior probability > 95 % under M8 was used to identify the positively selected sites.

## Author contribution

TK conceived and designed research strategy and wrote the paper. XY and YZ performed the experiments. XY, YZ and TK analyzed data.

## Acknowledgements

We thank Azusa Kadowaki for her help to run AlphaFold2 analysis.

