## Supplementary Figures for "Evolutionary history of metazoan TMEM16 family"

**Supplementary Figure 1. Molecular phylogenetic analysis of the transmembrane protein 16 (TMEM16) proteins across different metazoan species.**

The evolutionary history of metazoan TMEM16 proteins was based on the Multiple Sequence Comparison by Log-Expectation (MUSCLE) alignment and constructed on an IQ-Tree using LG+F+R10 as a model of amino acid substitution. The tree was rooted with eight human transmembrane channel-like proteins as an outgroup. The maximum-likelihood tree was based on 10,000 replicates of bootstrapping, and the branch support values (60–100) are indicated by the different sized brown circles. Scale bar represents the expected number of substitutions per site. The branches and protein names in the major subfamilies are colored, as shown in Figure 1.

**Supplementary Figure 2. Structures of anoctamin (ANO) domains in TMEM16A, F, and entire TMEM16K.**

Overall structures of ANO domains in murine TMEM16A (A) and F (B), as well as the whole human TMEM16K (C) predicted by AlphaFold2 are shown. Transmembrane (TM)1–10 domains, side chains of critical amino acid residues, and the main chain at the hinge region of the TM6 domain are colored, as shown in Figure 4. The distance between the two Ile residues in the TM4 and 6 domains to form the inner gate is also indicated. The Ile residue in the TM6 domain is substituted with the Leu residue in TMEM16K.

**Supplementary Figure 3. ANO domain structures of TMEM16A/Bs in various species.**

Overall structures of ANO domains in *Actinia equina* TMEM16A/B (A), *Nematostella vectensis* TMEM16A/B (B), *Clytia hemisphaerica* TMEM16A/B (C), *Lottia gigantea* XP.009060312.1 (D), *Capitella teleta* CapteP153854 (E) and CapteP153860 (F), *L. gigantea* XP.009065298.1 (G), *C. teleta* CapteP222303 (H), *Tribolium castaneum* XP.008192276.1 (I), *Drosophila melanogaster* NP.001245633.1 (CG10353, J), *T. castaneum* XP.008193545.1 (K), and *D. melanogaster* NP.001189248.1 (Subdued, L) predicted by AlphaFold2 are shown. Each structure is colored as shown in Figure 4.

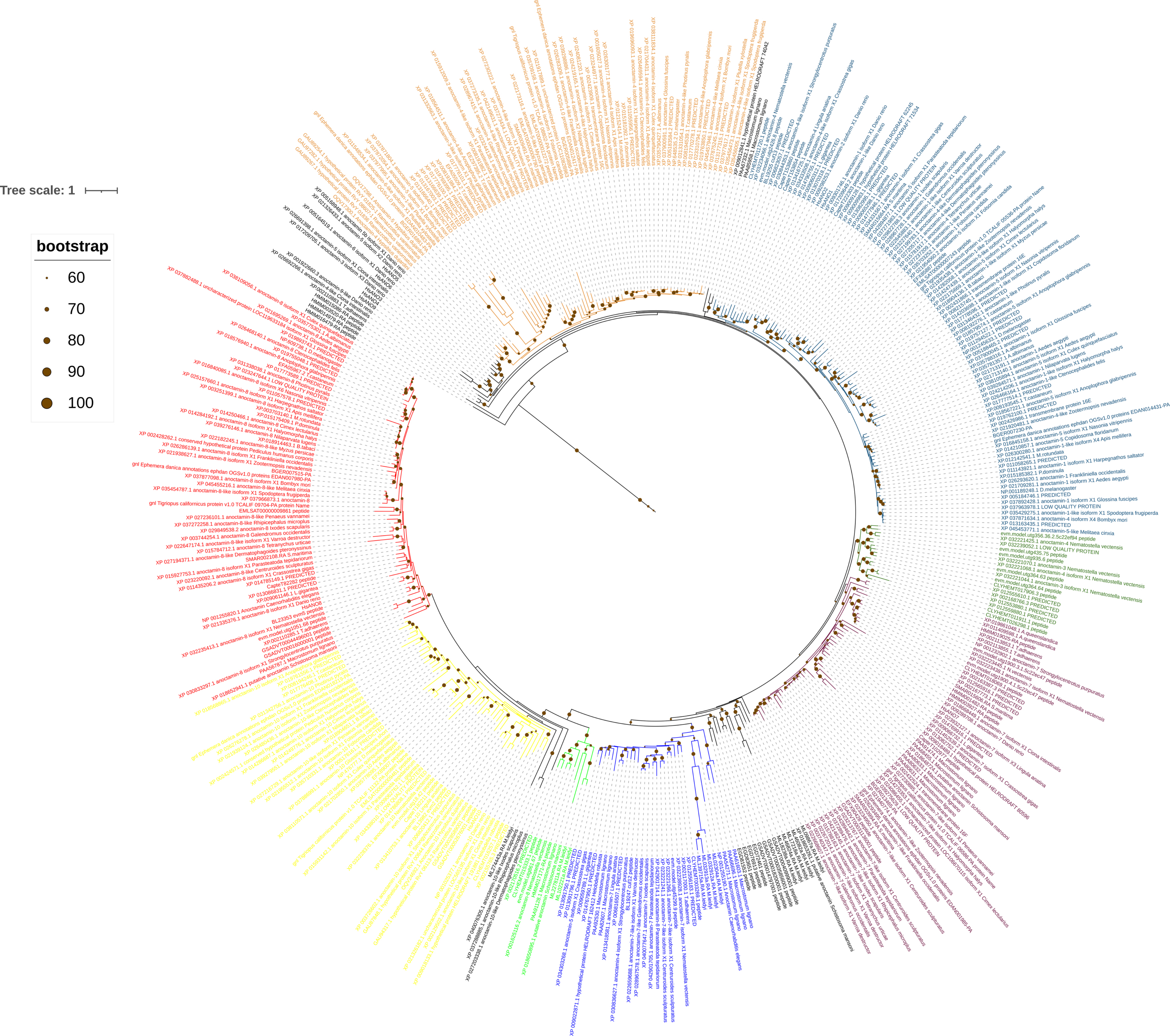
Tree scale: 1
bootstrap
60
70
80
90
100

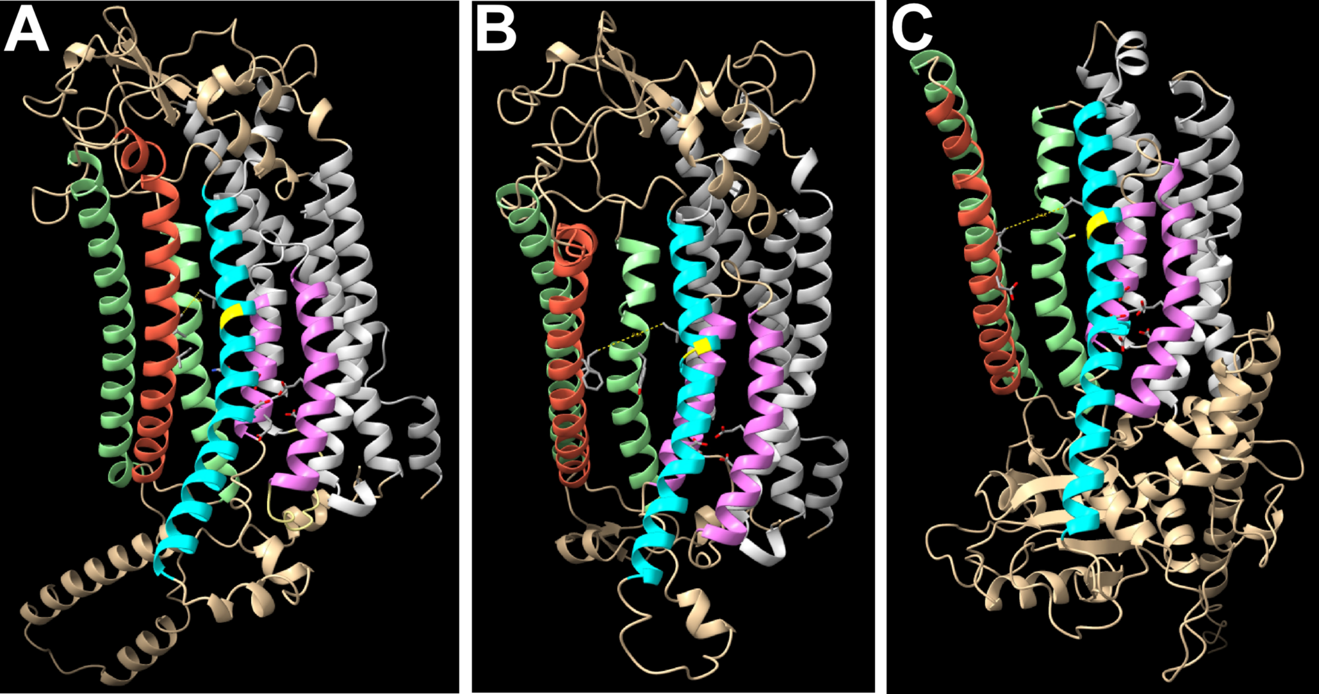
A
B
C

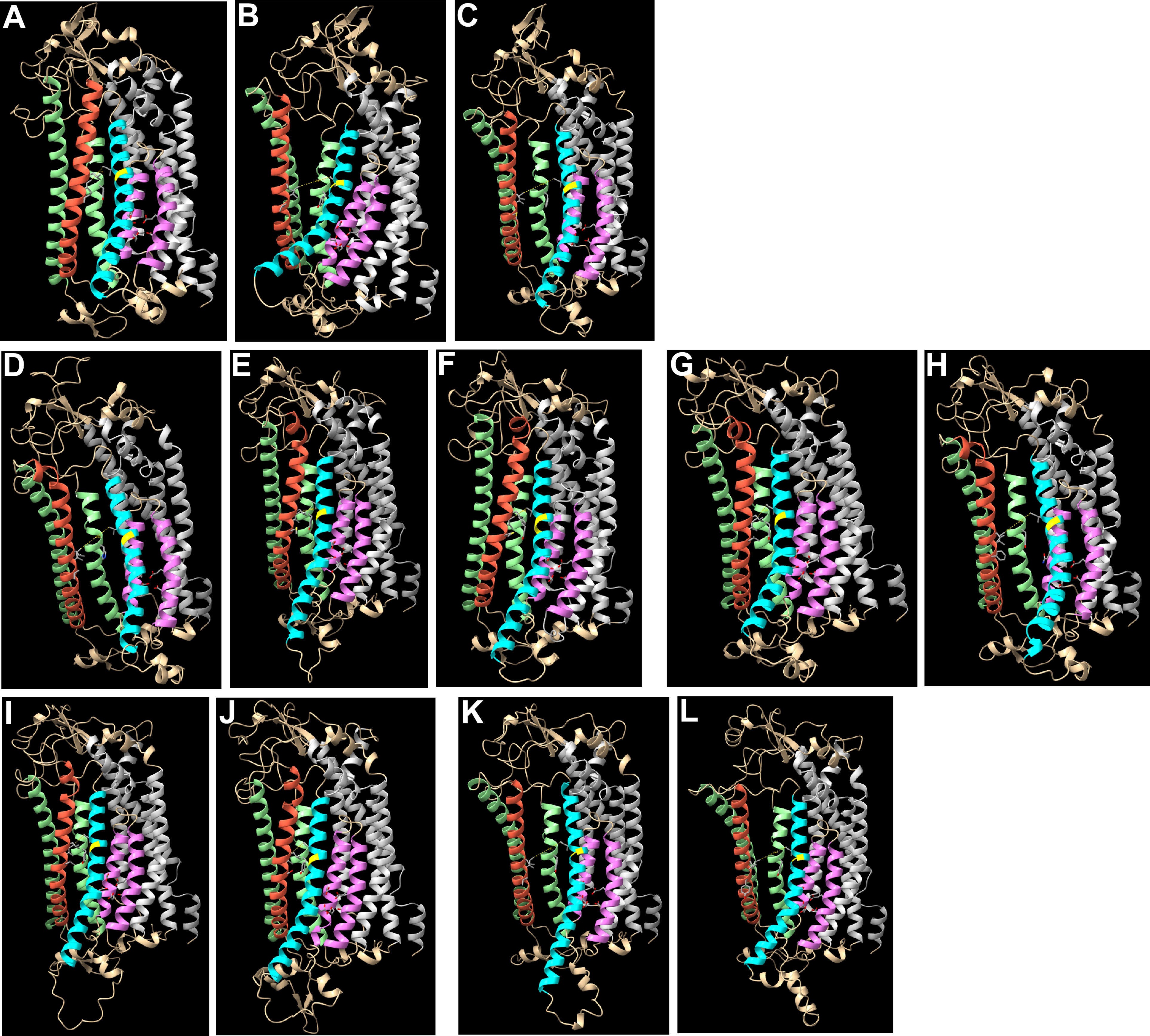
A
B
C
D
E
F
G
H
I
J
K
L
